# sPRR Signaling in Macrophages via the AT1R/Yap/Taz Axis to Induce Renal Fibrosis

**DOI:** 10.64898/2026.04.03.716436

**Authors:** Ye Feng, Huaqing Zheng, Shiying Xie, Fei Wang, Renfei Luo, Tianxin Yang

**Affiliations:** Department of Internal Medicine, University of Utah and Veterans Affairs Medical Center, Salt Lake City, Utah, USA

## Abstract

**Background:** Within the kidney, (pro) renin receptor (PRR) is abundantly expressed in the collecting duct (CD) and modulate physiological and pathophysiological processes. We have recently reported that activation of CD PRR mediates obstructive renal fibrosis in a mouse model of unilateral ureteral obstruction (UUO). The current study addresses the underlying mechanisms by examining the profibrotic pathway mediated by soluble PRR (sPRR)-dependent alternative macrophage activation.

**Methods:** We performed UUO or sham surgery on mice with CD-specific deletion of PRR (CD PRR KO) and floxed controls. After 7 days, we assessed fibrosis-related parameters, inflammatory cytokines, M1/M2 macrophage markers, other gene expression markers of kidney injury, and the concentration of plasma sPRR. We also administered vehicle or site-1 protease (S1P) inhibitor PF-429242 (PF) to C57BL/6 mice with UUO to determine the role of sPRR. Experiments were performed in vitro to examine the mechanism of sPRR-His-mediated macrophage M2 polarization and activation of potential target genes in bone-marrow-derived macrophages (BMDMs).

**Results:** Compared with the floxed control, CD PRR KO decreased macrophage accumulation, M2 polarization, and Yap/Taz expression while improving renal fibrosis and suppressing plasma sPRR levels following UUO. In BMDMs, sPRR-His treatment promoted macrophage M2 polarization, fibrosis, and Yap/Taz expression, which was dependent on angiotensin type 1 receptor (AT1R).

**Conclusion:** CD PRR-derived sPRR acts via ATR to promote macrophage M2 polarization and stimulates the AT1R/Yap/Taz axis, which leads to renal fibrosis during UUO.

## Introduction

Chronic kidney disease (CKD) involves gradual loss of kidney function (glomerular filtration rate (GFR))^1^ due to structural or functional anomalies of the kidneys. Regardless of etiology, the final common pathway of CKD is kidney fibrosis.^2,3^ The course of kidney fibrosis includes phases of priming, activation, execution, and progression.^4^ Myeloid cells accumulate in the glomerulus or tubulointerstitium, which is closely correlated with the progression of kidney fibrosis.^5^

Among inflammatory myeloid cells, both infiltrating and resident macrophages have profound effects on renal damage, repair, and fibrosis depending on their state of polarization.^6,7^ They may be activated and categorized as proinflammatory M1 macrophages that drive parenchymal injury via the generation of proinflammatory cytokines. However, they may also become alternatively activated as wound-healing or profibrotic M2 macrophages that facilitate tissue repair and tissue fibrosis. This effect occurs via the expression of immunosuppressive cytokines, including mannose receptor (MR), arginase-1 (Arg-1), resistin-like-α (Fizz1), and chitinase3 (YM1).^8,9^

(Pro)renin receptor (PRR) is a component of the renin-angiotensin system and are expressed in a wide variety of organs, including the kidneys,^10,11^ where PRR is predominantly expressed in the intercalated cells of the collecting ducts (CDs). Full-length PRR (fPRR) is a 350-amino-acid transmembrane protein that contains furin and site-1 protease (S1P) cleavage sites. S1P is a predominant protease that is responsible for the initial cleavage and release of a 28 kDa soluble PRR (sPRR).^12,13^ Although not required for the release of sPRR, furin-mediated second cleavage gives sPRR the ability to stimulate ENaC activity.^14^

Abundant evidence suggests that activation of PRR/sPRR plays an important role in the regulation of physiological and pathophysiological processes in the kidneys.^15–18^ In particular, our recent study indicates that the activation of renal PRR/sPRR plays a key role in mediating obstruction-induced renal fibrosis.^19^ However, the detailed signaling mechanisms behind this phenomenon remain elusive.

Hippo signaling is evolutionarily conserved and pivotal in regulating embryonic development and disease progression.^20^ Yes-associated protein (Yap) and transcriptional co-activator with PDZ-binding motif (Taz) serve as downstream effectors of this pathway. Yap/Taz can accumulate and relocate to the nucleus and interact with a number of transcription factors that may promote cell growth, differentiation, and survival after dephosphorylation.^21^ Blocking Yap/Taz can prevent tumor-associated macrophage M2 polarization.^22^

The signaling protein Wnt5a promotes TGFβ1-mediated macrophage polarization and kidney fibrosis by inducing transcriptional regulators Yap/Taz.^23^ Additionally, the Yap pathway is an important signaling branch downstream of angiotensin II type 1 receptor (AT1R) in cardiac fibroblast mechanotransduction.^24^ sPRR can directly interact with AT1R via lysine^199^ and asparagine.^295^ Therefore, the current study examined whether CD PRR/sPRR promotes macrophage M2 polarization and contributes to kidney fibrosis by sPRR-dependent activation of Yap/Taz activity.

## Methods

### Animal studies

Details about the generation of collecting duct-specific PRR knockout mice have been reported previously.^43^ The *PRR* gene is on the X-chromosome, so male mice heterozygous for floxed PRR and for AQP2-Cre (termed CD PRR KO) presumably have complete deletion of PRR in the CD. Only male CD PRR KO mice were used in the present study. Male 12-week-old C57BL/6 mice weighing approximately 18-20 g were purchased from Jackson Laboratory. All animals were housed in cages and maintained in a temperature-controlled room with a 12:12-hour light-dark cycle with free access to tap water and standard rat chow. All animal studies were conducted with the approval of the University of Utah Animal Care and Use Committee in accordance with the National Institutes of Health Guide for the Care and Use of Laboratory Animals.

Male 12-week-old CD PRR KO mice and floxed control mice from the same litter were subjected to the UUO procedure. Under general anesthesia with 2% isoflurane, a midline abdominal incision was made, and the left ureter was exposed and occluded by tightening the tubing with a 5-0 silk suture at the midportion. Sham-operated mice without ligating the ureter served as a control. For the pharmacological experiment, C57BL/6 mice underwent the same surgical procedure and were subcutaneously implanted with an osmotic minipump infusing vehicle or PF-429242 (20 mg/kg/d, Cat: HY-13447A, Medchem Express). Seven days after the surgery, tissues and blood samples were harvested under the general anesthesia.

### Western blot

Bone-marrow-derived macrophages (BMDMs) were lysed in 1× SDS sample buffer. The kidney tissues were lysed and then sonicated in PBS containing 1% Triton x-100, 250 μM phenylmethanesulfonyl fluoride (PMSF), 2 mM EDTA, and 5 mM dithiothreitol (DTT) (pH 7.5). The protein concentration was determined by a bicinchoninic acid (BCA) protein assay kit (Pierce Thermo-Scientific, Rockford, IL) according to the manufacturer’s instructions. An equal amount of protein was subjected to SDS-PAGE and transferred onto polyvinylidene difluoride (PVDF) membranes. The blots were blocked overnight with 5% nonfat dry milk in Tris-buffered saline (TBS), followed by incubation overnight with primary antibody.

After washing with TBS, the blots were incubated with horseradish peroxidase (HRP)-conjugated secondary antibody and visualized using Enhanced Chemiluminescence (ECL). Western blot was performed using anti-PRR antibody (cat: HPA003156, Sigma, 1:1000), anti-MR (cat: ab64693, Abcam, 1:1000), anti-β-Actin (cat: sc47778, Santa Cruz Biotechnology, USA, 1:1000), anti-p-Yap (Ser127) (cat: 13008S, Cell Signaling Technology, 1:1000), anti-Yap/Taz (cat: 8418, Cell Signaling Technology, 1:1000), and anti-TEAD (cat: 2530, Cell Signaling Technology, 1:1000). Quantification was performed by measuring the intensity of the signals using the Image J software package (National Institutes of Health).

### qRT-PCR

Snap-frozen renal samples were homogenized in TRIzol reagent (Cat: 15596018, Life Technologies, CA, USA). Total RNA isolation and reverse transcription (RT) were performed according to the manufacturer’s recommendations. We used 1 μg of total RNA as a template for RT and a Transcriptor First Strand cDNA Synthesis Kit (Cat: 4368813, Thermo Fisher) according to the manufacturer’s instructions. qPCR was performed using the ABI Prism StepOnePlus System (Applied Biosystems, Life Technologies, USA) and the SYBR Premix Ex Taq kit (Tli RNaseH Plus) (Cat: 1803132, Thermo Fisher) according to the manufacturer’s instructions. The primers are listed in Table S1. The amount of mRNA or gene relative to the internal control was calculated as 2^(−Δ*CT)*, where *ΔCT = CT_gene_ - CT_control_*.

### Histology and Immunohistochemical Staining

Kidney samples from mice were fixed in 10% neutral formalin and embedded in paraffin. 3-μm-thick sections were stained with periodic acid-Schiff and Masson stain. Images were captured using a Leica DMI4000B fluorescence microscope (Wetzlar, Germany).

### Immunofluorescence Staining

Kidney cryosections of 3-μm thickness were fixed for 15 minutes with 4% paraformaldehyde followed by permeabilization with 0.2% Triton X-100 in 1× PBS for 5 minutes at room temperature. After blocking with 2% donkey serum for 60 minutes, the slides were immunostained with the following antibodies: anti-F4/80 (cat: 14-4801, eBioscience, San Diego, CA), anti-MR (cat: ab64693, Abcam), and anti-Yap/Taz (cat: 8418, Cell Signaling Technology). BMDMs cultured on coverslips were washed with cold 1× PBS and fixed with cold methanol and acetone (1:1) for 10 minutes at −20°C. After three extensive washings with 1× PBS, the cells were treated with 0.1% Triton X-100 for 5 minutes, blocked with 2% normal donkey serum in 1× PBS buffer for 40 minutes at room temperature, and incubated with anti-Yap/Taz (cat: 8418, Cell Signaling Technology). Cells were also stained with DAPI to visualize the nuclei. Images were captured using a Leica DMI4000B fluorescence microscope (Wetzlar, Germany).

### Cell Culture and Treatment

BMDMs were isolated as described previously.^25^ BMDMs were cultured in DMEM containing 10% (vol/vol) FBS, 10 ng/ml mouse M-CSF (cat: 416-ML-050; R&D Systems, Minneapolis, MN), and 1% (vol/vol) antibiotics for 9 days. The medium was changed every other day. BMDMs were cultured with serum-free medium and treated with sPRR-His (10 nM, Xbio, Shanghai, China) for various durations. The BMDMs were treated with losartan (10 μM, cat: HY-17512; Medchem express), verteporfin (10 nM, cat: SML0534; Sigma-Aldrich), or PF-429242 (10 μM, cat: HY-13447A, Medchem Express) for 1 hour to block AT1, Yap, and S1P, respectively. This was followed by the addition of sPRR-His or TGFβ1 (2ng/ml, cat: 100-B-010-CF; R&D Systems) for 24 hours.

### Quantitative Analyses of Total Collagen in Kidney Tissue

The total collagen within kidney tissue was quantitated as reported previously.^25^ Briefly, 3-μm sections of paraffin-embedded tissue were stained overnight with Sirius red F3BA and Fast Green FCF (Sigma-Aldrich). After washing three times with 1× PBS buffer, the dye was eluted from tissue sections with 0.1 N sodium hydroxide methanol. Absorbance values at 540 and 605 nm were determined for Sirius red F3BA and Fast Green FCF binding protein, respectively. Measurements of the ratio of collagen to total protein were expressed in micrograms per milligram of total protein.

### Monocyte Macrophage Isolation and Enzyme Immunoassay

The kidneys of mice were harvested, minced, and incubated with 1 mg/ml collagenase (cat: 17018-017; Gibco) and 0.1 mg/ml DNase (cat: 10104159001; Roche) for 30 minutes at 37°C. The spleens were also harvested, and minced, followed by incubation with 0.1 mg/ml DNAase for 30 minutes at 37°C. Single-cell suspensions were obtained by filtering through a 40-μm cell strainer and incubated with PE-CD115 antibody (cat: AFS98; Biolegend). Macrophages were then enriched from the single-cell suspensions with a mouse PE positive selection kit II (cat: 17666; Easy Sep) according to the manufacturer’s instructions. The amount of sPRR in plasma was determined by using commercially available enzyme immunoassay (EIA) kits for sPRR according to the manufacturer’s instructions (cat no. JP27782, IBL).

### Statistics

Data are summarized as the mean ± the standard error of the mean. All data points represent animals that were included in the statistical analyses. Sample sizes were determined based on similar previous studies or pilot experiments. Statistical analysis for animal experiments was performed using analysis of variance (ANOVA) with the Bonferroni test for multiple comparisons or by two-tailed paired or unpaired Student’s *t* tests for two comparisons. Values of *p* < 0.05 were considered statistically significant.

## Results

### Ablation of PRR in CD Attenuates Macrophage Accumulation and M2 Polarization in UUO Kidneys

We previously reported a novel profibrotic role of CD PRR/sPRR in a mouse model of UUO.^19^ As shown in Supplemental Figures 1 and 2, CD PRR KO significantly ameliorated UUO-induced renal fibrosis. The present study examined the underlying mechanisms with emphasis on macrophage polarization. We subjected CD PRR^f/f^ mice and CD PRR KO controls to 7-day UUO and analyzed the status of macrophages. By immunofluorescence using an antibody against F4/80, macrophage accumulation was remarkably increased in the obstructed kidneys from CD PRR^f/f^ mice, whereas the increase was much less in those from CD PRR KO mice (Figure 1A). We sorted macrophages from the spleens and obstructed kidneys of the two genotypes and examined the abundance for mannose receptor (MR), a marker for M2 polarized macrophages. The western blot results showed that the protein expression of MR was significantly decreased in macrophages isolated from obstructed kidneys from CD PRR KO mice compared to those from CD PRR^f/f^ mice (Figure 1B). In contrast MR protein expression was unaffected by UUO or the genotype (Figure 1B). Moreover, immunofluorescence staining demonstrated that MR expression was decreased in macrophages from the obstructed kidneys from CD PRR KO mice compared to those from CD PRR^f/f^ mice (Figure 1C).

**Figure 1.**
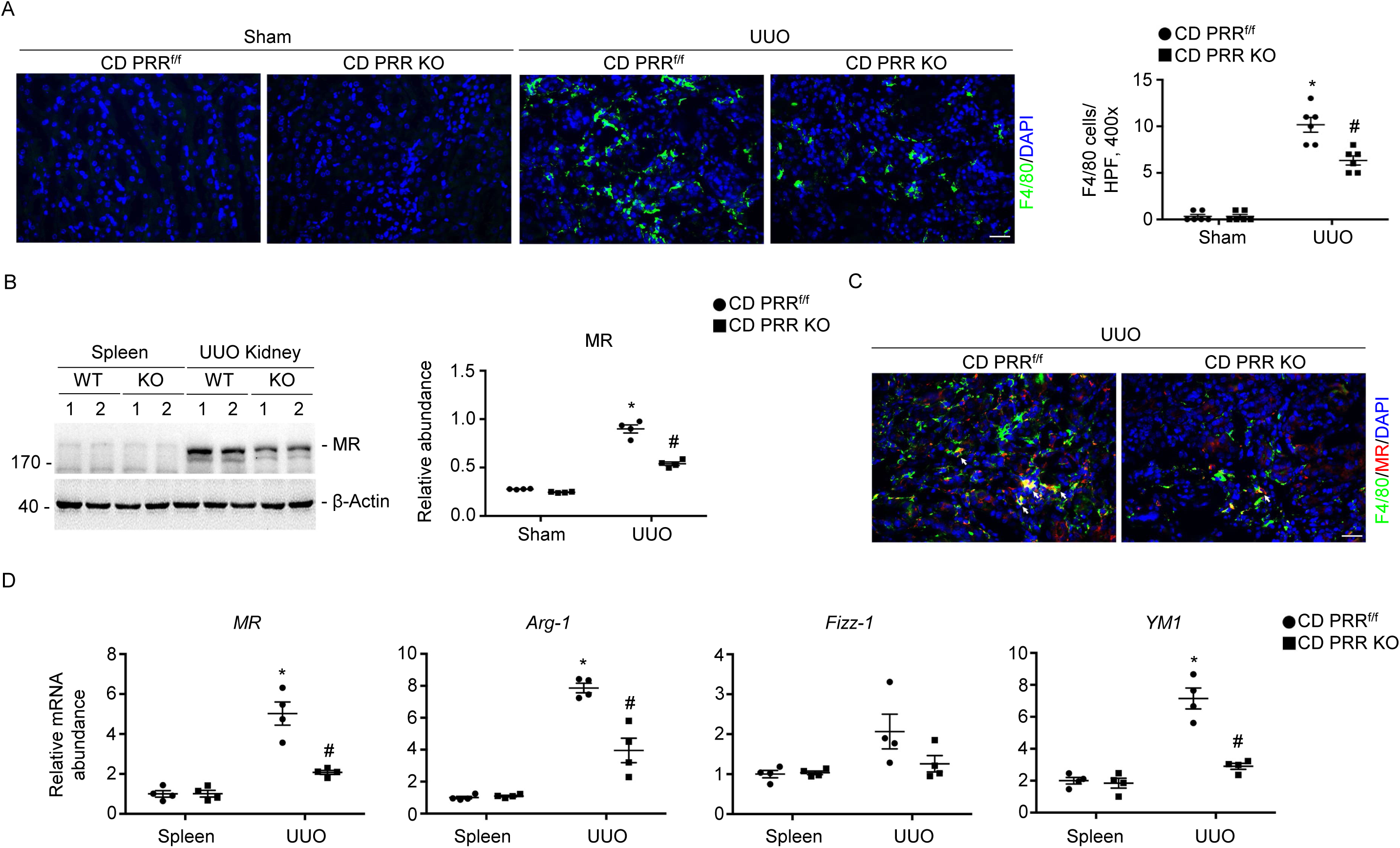
Ablation of PRR in the CD attenuates macrophage accumulation and M2 polarization in obstructed kidneys. (A) Representative micrographs (left) and quantitative analysis (right) of F4/80-positive cells in the kidneys of male 12-week-old CD PRR^f/f^ and CD PRR KO mice subjected to sham operation or UUO. Scale bar = 50 μm. * *p* < 0.05 versus CD PRR^f/f^ sham control, *n* = 6; ^#^ *p* < 0.05 versus CD PRR^f/f^ littermates after UUO, *n* = 6. (B) Western blot analyses (left) and quantitative analysis (right) of MR in macrophages sorted from spleen and obstructed kidneys with CD115 microbeads. Numbers (1, 2) indicate individual sample that was pooled from three individual animals within the same group. * *p* < 0.05 versus spleens from CD PRR^f/f^ littermates after UUO, *n =* 4; ^#^ *p* < 0.05 versus UUO kidneys from CD PRR^f/f^ littermates after UUO, *n =* 4. (C) Representative co-immunostaining images of F4/80 and MR in the UUO kidneys among indicated groups. White arrows indicate co-staining positive cells. Scale bar = 50 μm. (D) qRT-PCR analysis of mRNA expression of *Arg-1*, *MR*, *Fizz-1,* and *YM1* (left to right in order) in macrophages isolated from spleen and obstructed kidneys with CD115 microbeads at 7 days after surgery. Each sample was pooled from three individual animals within the same group. * *p* < 0.05 versus spleens from CD PRR^f/f^ littermates after UUO, *n =* 4; ^#^ *p* < 0.05 versus UUO kidneys from CD PRR^f/f^ littermates after UUO, *n =* 4.

By qRT-PCR, the mRNA abundance was significantly reduced for M2 macrophage markers in macrophages from the obstructed kidneys of CD PRR KO mice compared to those from CD PRR^f/f^ mice, including Arg-1, MR, Fizz-1, and YM1 (Figure 1D). These results suggest that conditional deletion of PRR in the CD may attenuate macrophage M2 polarization, thus mitigating renal fibrosis.

### Ablation of PRR in CD Mitigates Macrophage Yap/Taz Expression in Fibrotic Kidneys

We next investigated the molecular mechanisms of CD PRR-dependent macrophage M2 polarization in the UUO model. Given the recent report of the novel role of the Yap/Taz activation in mediating macrophage M2 polarization during kidney fibrosis,^23^ we examine the status of the Yap/Taz axis in macrophages from the obstructed kidneys. Western blot analysis showed that Yap/Taz expression was significantly increased in macrophages isolated from the obstructed kidneys of CD PRR^f/f^ mice, and this increase was much less in those of CD PRR KO mice (Figure 2A). Consistent with this observation, immunofluorescence staining showed a reduction of Yap/Taz expression in macrophages from obstructed kidneys of CD PRR KO mice compared to those from CD PRR^f/f^ mice (Figure 2B). A similar pattern of changes between the two genotypes occurred in the mRNA expression of Yap, Taz, and their target genes, including connective tissue growth factor (*Ctgf*) and ankyrin repeat domain 1 (*Ankrd1*) in macrophages from obstructed kidneys (Figure 2C). Therefore, these results suggest that deletion of CD PRR downregulated macrophage Yap/Taz expression, resulting in reduced macrophage M2 polarization and thus improved renal fibrosis during UUO.

**Figure 2.**
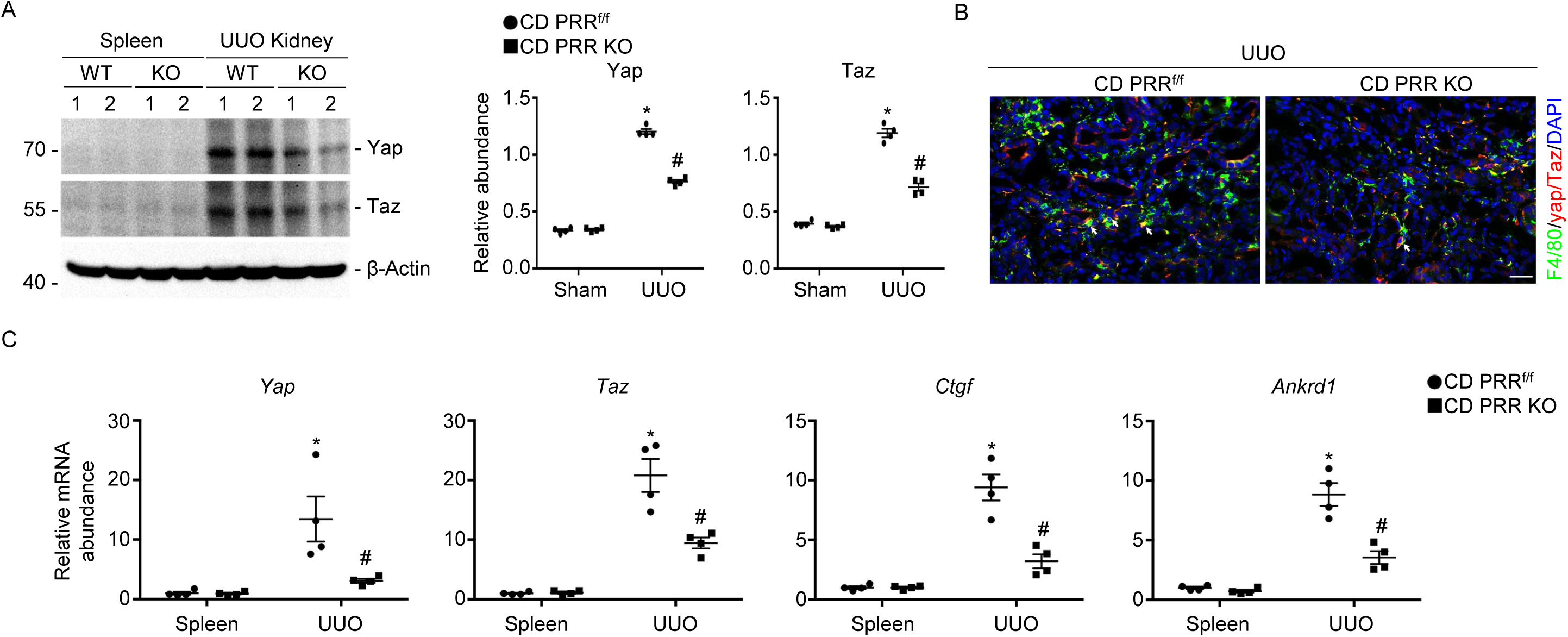
Ablation of PRR in the CD mitigates macrophage Yap/Taz expression in obstructed kidneys. (A) Representative Western blots (left) and densitometric analysis (right) of Yap/Taz in macrophages isolated from spleen and obstructed kidneys using CD115 microbeads. Numbers (1, 2) indicate individual sample that was pooled from three individual animals within the same group. \**p <* 0.05 versus spleens from CD PRR^f/f^ littermates after UUO, *n =* 4; ^#^*p <* 0.05 versus UUO kidneys from CD PRR^f/f^ littermates after UUO, *n =* 4. (B) Representative co-immunostaining images of F4/80 and Yap/Taz in UUO kidneys among indicated groups. White arrows indicate co-staining positive macrophages. Scale bar = 50 μm. (C) qRT-PCR analysis of mRNA expression of *Yap, Taz, Ctgf*, and *Ankrd1* (left to right in order) in macrophages isolated from spleen and the obstructed kidneys using CD115 microbeads. Each sample was pooled from three individual animals within the same group. \**p <* 0.05 versus spleens from CD PRR^f/f^ littermates after UUO, *n =* 4; ^#^*p <* 0.05 versus UUO kidneys from CD PRR^f/f^ littermates after UUO, *n =* 4.

### CD PRR Promotes Macrophage M2 Polarization and Yap/Taz Expression via S1P- Derived sPRR

sPRR is predominantly generated by site-1 protease (S1P)-mediated cleavage of full-length PRR.^26^ Emerging evidence demonstrates that S1P-derived sPRR mediates multiple physiological and pathophysiological process, particularly renal fibrosis.^27^ We examined whether inhibition of S1P-derived sPRR mediates renal fibrosis through the activation of macrophage M2 polarization and Yap/Taz axis. We administered the S1P inhibitor PF-429242 or vehicle to UUO mice for 7 days and then examined the status of macrophage M2 polarization and Yap/Taz axis. As shown in Figure 3B, macrophage accumulation was remarkably decreased in the obstructed kidneys from mice infused with PF compared to the vehicle control.

**Figure 3.**
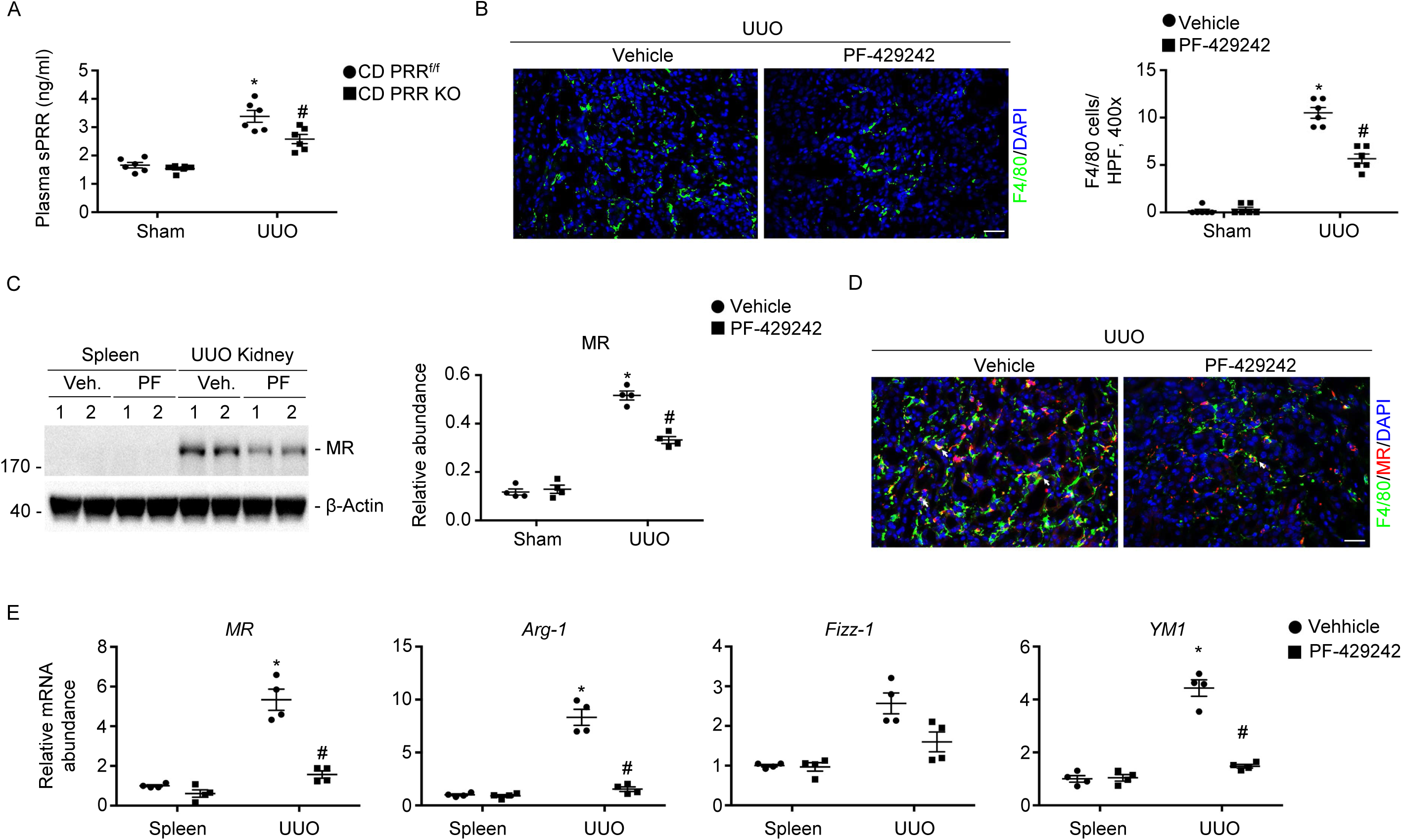
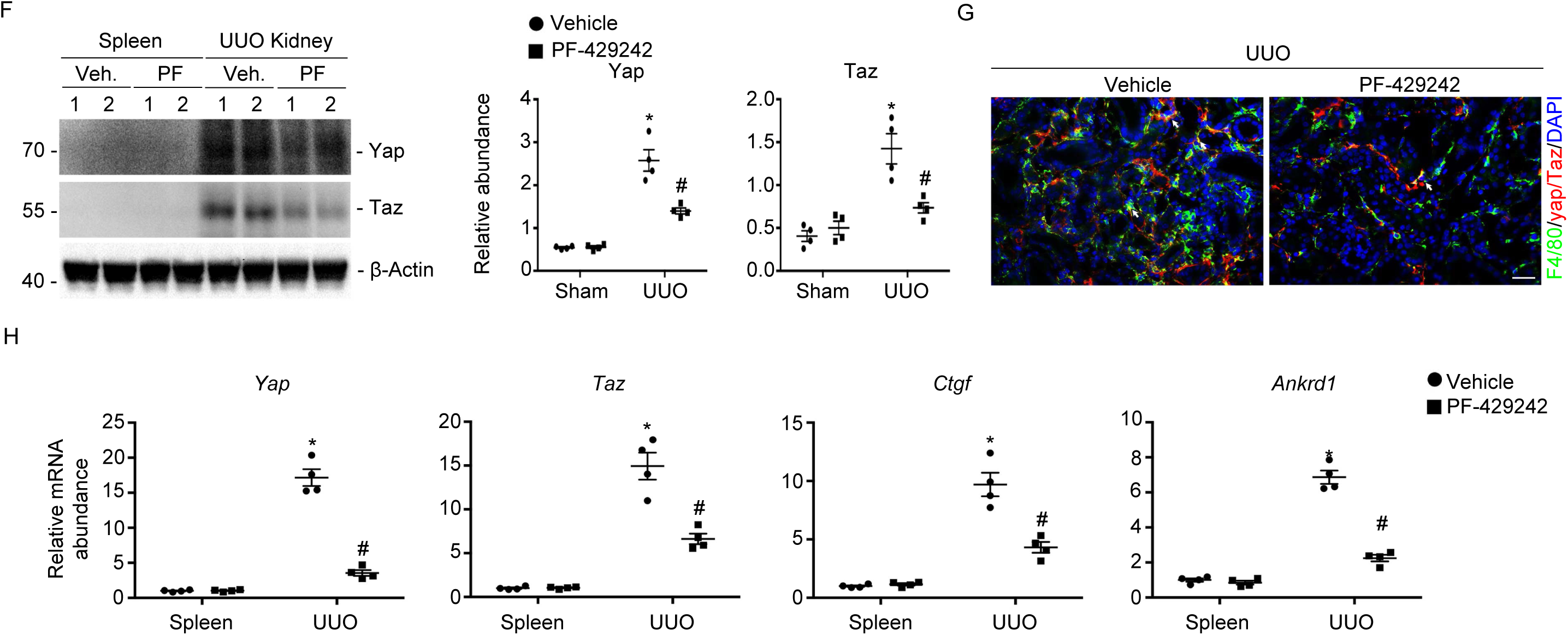
Blockade of sPRR production mitigates macrophage accumulation and M2 polarization in obstructed kidneys. (A) Plasma samples from CD PRR^f/f^ and CD PRR KO mice were subjected to ELISA measurement of plasma sPRR concentration. \**p <* 0.05 versus CD PRR^f/f^ sham control, *n =* 6; ^#^*p <* 0.05 CD PRR^f/f^ littermates after UUO, *n =* 6. (B) Male 12-week-old C57/Representative micrographs (left) and quantitative analysis (right) of F4/80-positive cells in kidneys. Scale bar = 50 μm. \**p <* 0.05 versus control kidneys from mice infused with vehicle, *n =* 6; ^#^*P*<0.05 versus obstructed kidneys from mice infused with vehicle, *n =* 6. (C) Western blot analyses (left) and quantitative analysis (right) of MR in macrophages sorted from spleen and UUO kidneys with CD115 microbeads. Numbers (1, 2) indicate individual sample that was pooled from three individual animals within the same group. \**p <* 0.05 versus spleens from mice infused with vehicle, *n =* 4; ^#^*p <* 0.05 versus UUO kidneys from vehicle littermates after UUO, *n =* 4. (D) Representative co-immunostaining images of F4/80 and MR in the UUO kidneys among indicated groups. White arrows indicate co-staining positive cells. Scale bar = 50 μm. (E) Real-time qRT-PCR analysis showing the mRNA abundance of *Arg-1*, *MR*, *Fizz-1,* and *YM1* in macrophages enriched from spleen and UUO kidneys with CD115 microbeads at 7 days after surgery. Each sample was pooled from three individual animals within the same group. \**p <* 0.05 versus spleens from mice infused with vehicle after UUO, *n =* 4; ^#^*p <* 0.05 versus UUO kidneys from mice infused with vehicle, *n =* 4. (F) Western blot analyses (left) and quantitative analysis (right) of Yap/Taz in macrophages sorted from spleen and UUO kidneys with CD115 microbeads. Numbers (1, 2) indicate individual sample that was pooled from three individual animals within the same group. \**p <* 0.05 versus spleens from mice infused with vehicle, *n =* 4; ^#^*p <* 0.05 versus UUO kidneys from vehicle littermates after UUO, *n =* 4. (G) Representative co-immunostaining images of F4/80 and Yap/Taz in UUO kidneys among indicated groups. White arrows indicate co-staining positive macrophages. Scale bar = 50 μm. (H) Real-time qRT-PCR analysis showing the mRNA abundance of *Yap, Taz, Ctgf,* and *Ankrd1* in macrophages sorted from spleen and UUO kidneys with CD115 microbeads. Each sample was pooled from three individual animals within the same group. \**p <* 0.05 versus spleens from mice infused with vehicle after UUO, *n =* 4; ^#^*p <* 0.05 versus UUO kidneys from mice infused with vehicle, *n =* 4.

We also examined the MR expression in the same macrophage preparations. Western blot analysis showed that PF treatment reduced MR protein expression in macrophages of obstructed kidneys compared with the vehicle control, while there was a lack of MR expression in macrophages from the spleen irrespective of PF treatment (Figure 3C). In addition, immunofluorescence staining demonstrated that MR expression was decreased in macrophages from UUO/PF mice compared to the vehicle control (Figure 3D). In obstructed kidney-derived macrophages, PF treatment reduced the mRNA expression of M2 macrophage markers, including Arg-1, MR, Fizz-1, and YM1 (Figure 3E).

We examined the Yap/Taz expression in the same macrophage preparations. Western blot analysis showed that the expression in obstructed kidney-derived macrophages was significantly suppressed by PF treatment, while there was a lack of expression in spleen-derived macrophages (Figure 3F). Immunofluorescence staining showed a similar reduction of Yap/Taz expression in macrophages from obstructed kidneys following PF treatment (Figure 3G).

The same pattern of changes occurred in the mRNA expression of Yap, Taz, and target genes (including *Ctgf* and *Ankrd1*) as Yap/Taz protein expression (Figure 3H). These results suggest that S1P-derived sPRR drives macrophage M2 polarization and Yap/Taz expression, which contribute to obstruction-induced renal fibrosis.

### sPRR Directly Promotes Alternative Macrophage Activation

To explore the direct role of sPRR in regulating macrophage M2 polarization, we treated BMDMs with sPRR-His and examined the changes in the markers indicative of M2 polarization. According to Western blot analysis, 24-hour exposure to sPRR-His treatment at 10 nM significantly induced MR protein expression in BMDMs (Figure 4A). Consistent with this observation, the sPRR-His treatment induced parallel upregulation of the mRNA expression of MR, Arg-1, MR, Fizz-1, and YM1 (Figure 4B). Interestingly, mRNA abundance for M1 macrophage markers, including IL-6 and IL-12b, were decreased in macrophages induced by sPRR-His (Supplemental Figure 3). Moreover, PF inhibited macrophage M2 polarization induced by TGFβ in BMDMs (Supplemental Figure 4).

**Figure 4.**
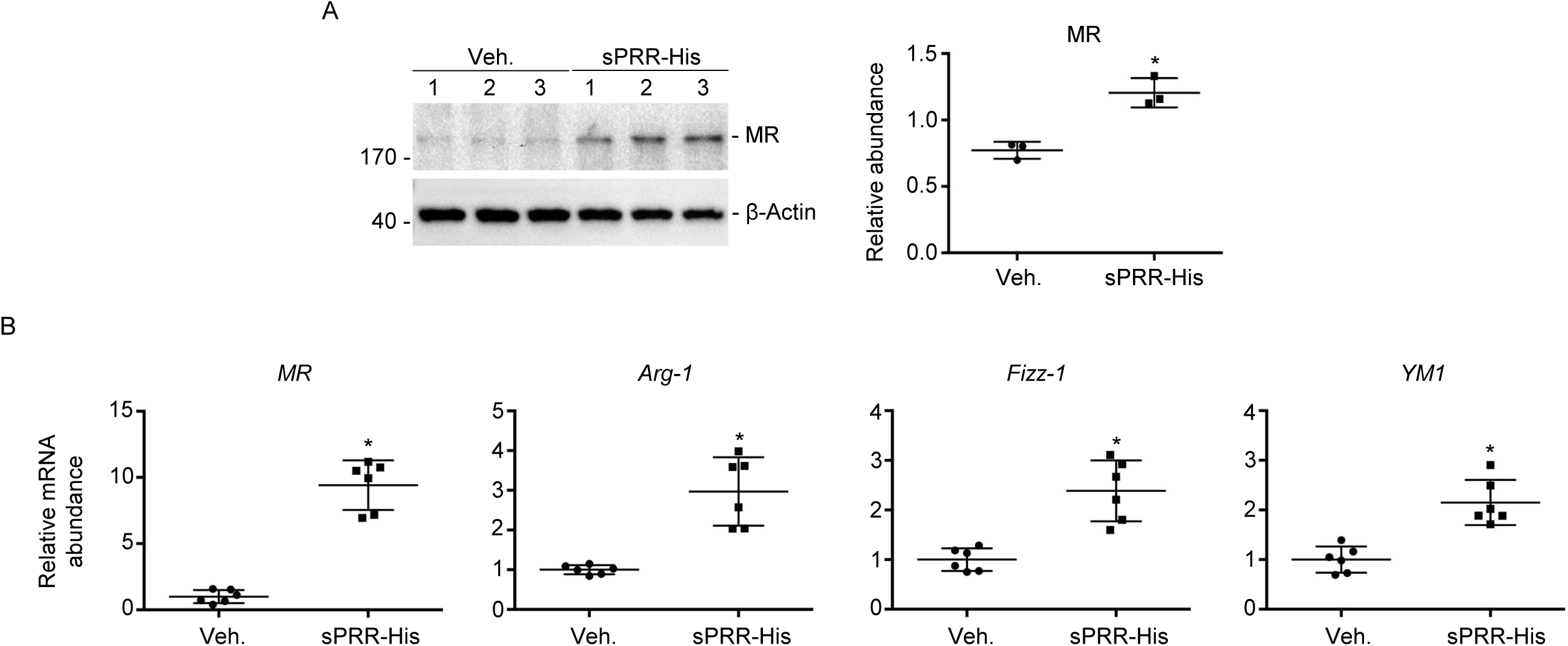
sPRR-His directly promotes macrophage alternative activation. (A) Western blot analyses (left) and quantitative analysis (right) of MR in BMDMs treated with 10 nM sPRR-His for 24 hours. Numbers (1–3) indicate individual sample within the same group. \**p <* 0.05 versus BMDMs treated with vehicle alone, *n =* 3. (B) qRT-PCR analysis showing the mRNA abundance of *Arg-1, MR, Fizz-1*, and *YM1* in BMDMs that were treated with 10 nM sPRR-His for 24 hours. qRT-PCR data were normalized to reference gene *Gapdh*. \**p <* 0.05 versus BMDMs treated with vehicle alone, *n =* 6.

We examined the effect of fibrosis in BMDMs exposed to sPRR-His. The expression of FN and α-SMA were significantly induced by sPRR-His treatment (Figure 5B). In parallel, the mRNA levels of TGFβ1, TGFβ3, PDGFA, PDGFB, PDGFC, PDGFD, VEGFA, and VEGFC in BMDMs were all elevated (Figure 5C). These results demonstrate that sPRR has a direct role in promoting macrophage M2 polarization and thus the fibrotic response in macrophages.

**Figure 5.**
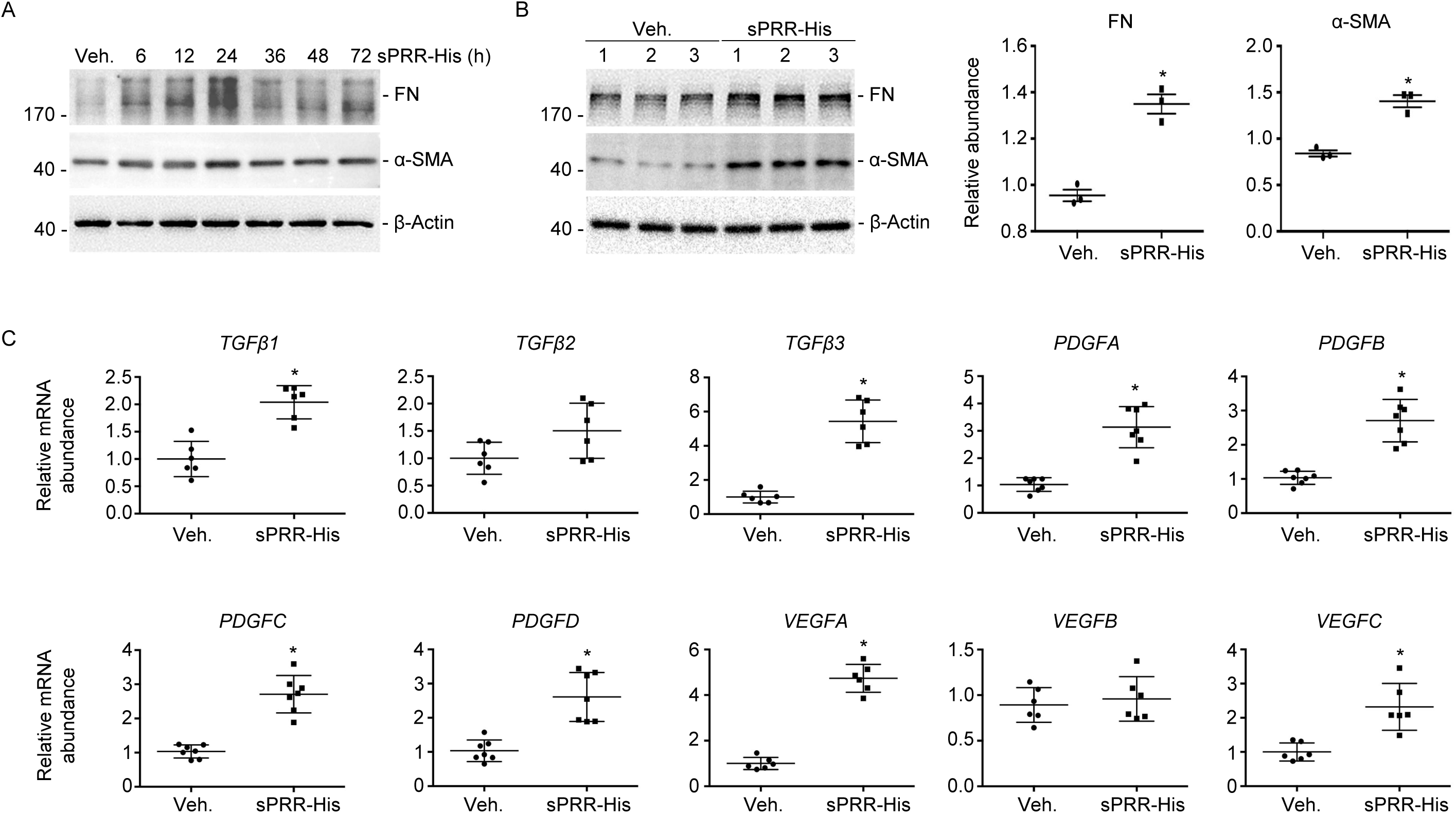
sPRR-His directly promotes fibrosis in BMDMs. (A) Western blot analyses showing the abundance of FN and αSMA in BMDMs treated with 10 nM sPRR-His for different indicated times. (B) Western blot analyses (left) and quantitative analysis (right) of FN and αSMA in BMDMs treated with 10 nM sPRR-His for 24 hours. Numbers (1–3) indicate individual sample within the same group. \**p <* 0.05 versus BMDMs treated with vehicle alone, *n =* 3. (C) Real-time qRT-PCR analysis showing mRNA abundance of *TGFβ1-3, PDGFA-D,* and *VEGFA-C* in BMDMs treated with 10 nM sPRR-His for 24 hours. qRT-PCR data were normalized to reference gene *Gapdh*. \**p <* 0.05 versus BMDMs treated with vehicle alone, *n =* 6.

### sPRR Signaling through AT1R Induces Macrophage M2 Polarization

Our recent study showed that sPRR induces obesity-related hypertension through endothelial dysfunction by directly interacting with AT1R.^28^ Therefore, we examined whether AT1R similarly mediates the action of sPRR in macrophages. Indeed, in a wide array of kidney diseases, AT1R is expressed in the immune cells such as macrophages to mediate kidney fibrosis.^29^ Thus, we tested whether sPRR-His-induced macrophage M2 polarization and fibrosis depend on AT1R.

BMDMs were treated with sPRR-His with or without losartan for 24 hours. As shown in Figure 6A, the band of ∼30 kDa detected by anti-PRR antibody was absent in the non-sPRR-His treated group, suggesting a low level of or no expression of endogenous sPRR in the cultured cells, but was abundant in sPRR-His treated group. This result may indicate the entry of sPRR-His into the cells. Interestingly, losartan remarkably decreased the sPRR-His content. Furthermore, it effectively suppressed sPRR-His-induced FN and the expressions of α-SMA, Arg-1, MR, Fizz-1, and YM1 in BMDMs (Figure 6B). The data suggests that sPRR-His induces macrophage M2 polarization and fibrosis via AT1R.

**Figure 6.**
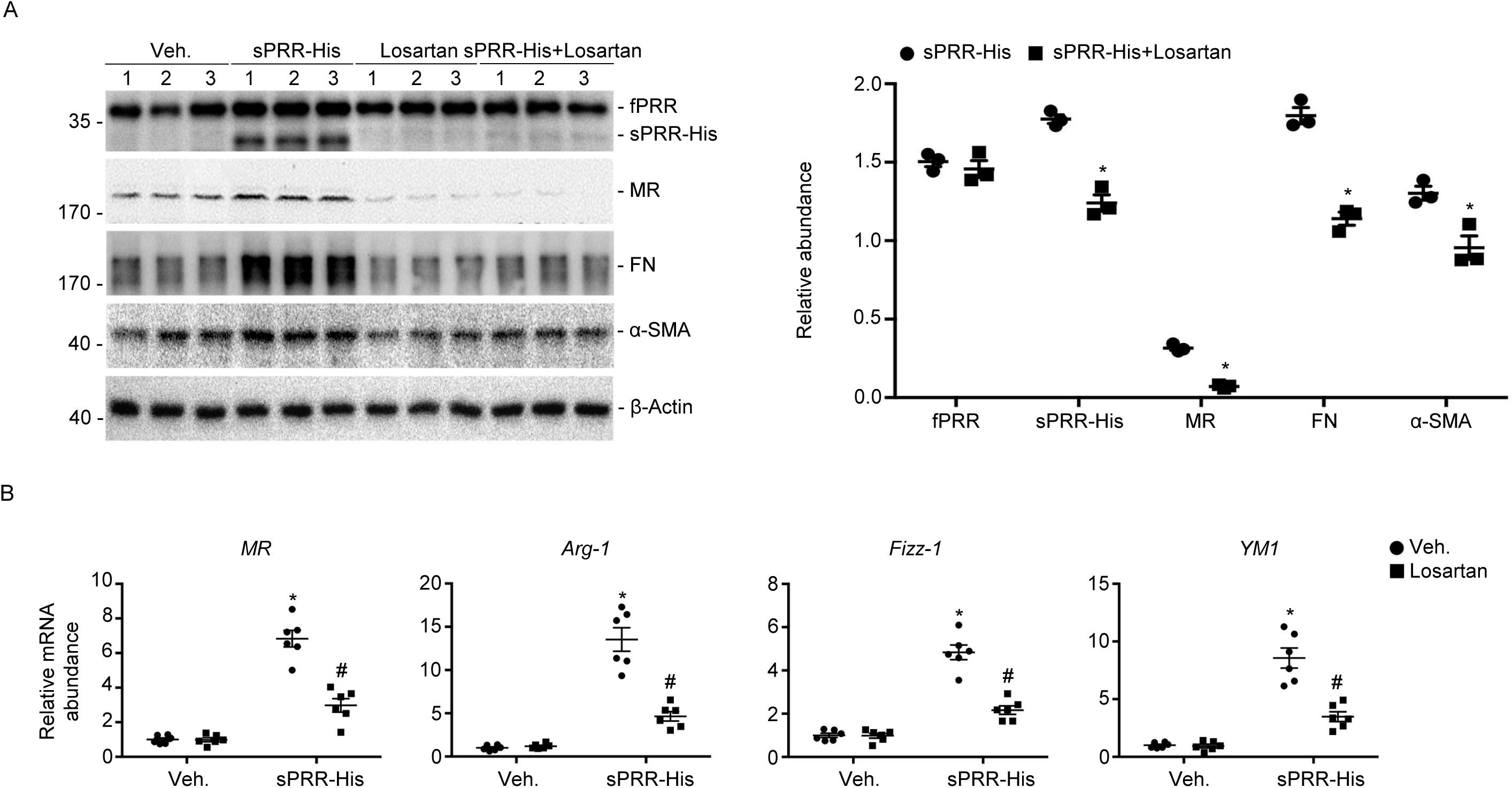
AT1 receptor inhibitor losartan blocked sPRR-His, induced alternative macrophage activation, and fibrosis in BMDMs. (A) Western blot analyses (left) and quantitative analysis (right) of fPRR, sPRR-His, MR, FN, and αSMA in BMDMs pretreated with 10 μM losartan of 1 h and then 10 nM sPRR-His for 24 hours. Numbers (1–3) indicate individual sample within the same group. \**p <* 0.05 versus BMDMs treated with sPRR-His, *n =* 3. (B) Real-time qRT-PCR analysis showing mRNA abundance of *Arg-1, MR, Fizz-1*, and *YM1* in BMDMs pretreated with 10 μM: Losartan for 1 hour and then 10 nM sPRR-His for 24 hours. qRT-PCR data were normalized to reference gene *Gapdh*. \**p <* 0.05 versus BMDMs treated with vehicle alone, *n =* 6; ^#^*p <* 0.05 versus BMDMs treated with sPRR-His, *n =* 6.

### Yap/Taz Mediates sPRR-His-Induced Macrophage Alternative Activation and Fibrosis

The YAP pathway is an important signaling branch that is downstream of AT1R in cardiac fibroblast mechanotransduction.^24^ Furthermore, Yap/Taz mediates Wnt5a-exacerbated macrophage M2 polarization and kidney fibrosis.^23^ We investigated whether sPRR-His induces macrophage M2 polarization and fibrosis through Yap/Taz activation by testing Yap/Taz expression in BMDMs after sPRR-His treatment. As shown in Figure 7A, the expression was increased in BMDMs after sPRR-His treatment, and sPRR-His promoted nuclear translocation of Yap/Taz in BMDMs, which could be blocked by losartan (Figure 7B). PF largely inhibited the TGFβ1-induced mRNA abundance of Yap, Taz, and target genes including *Ctgf* and *Ankrd1* in BMDMs (Supplemental Figure 5).

**Figure 7.**
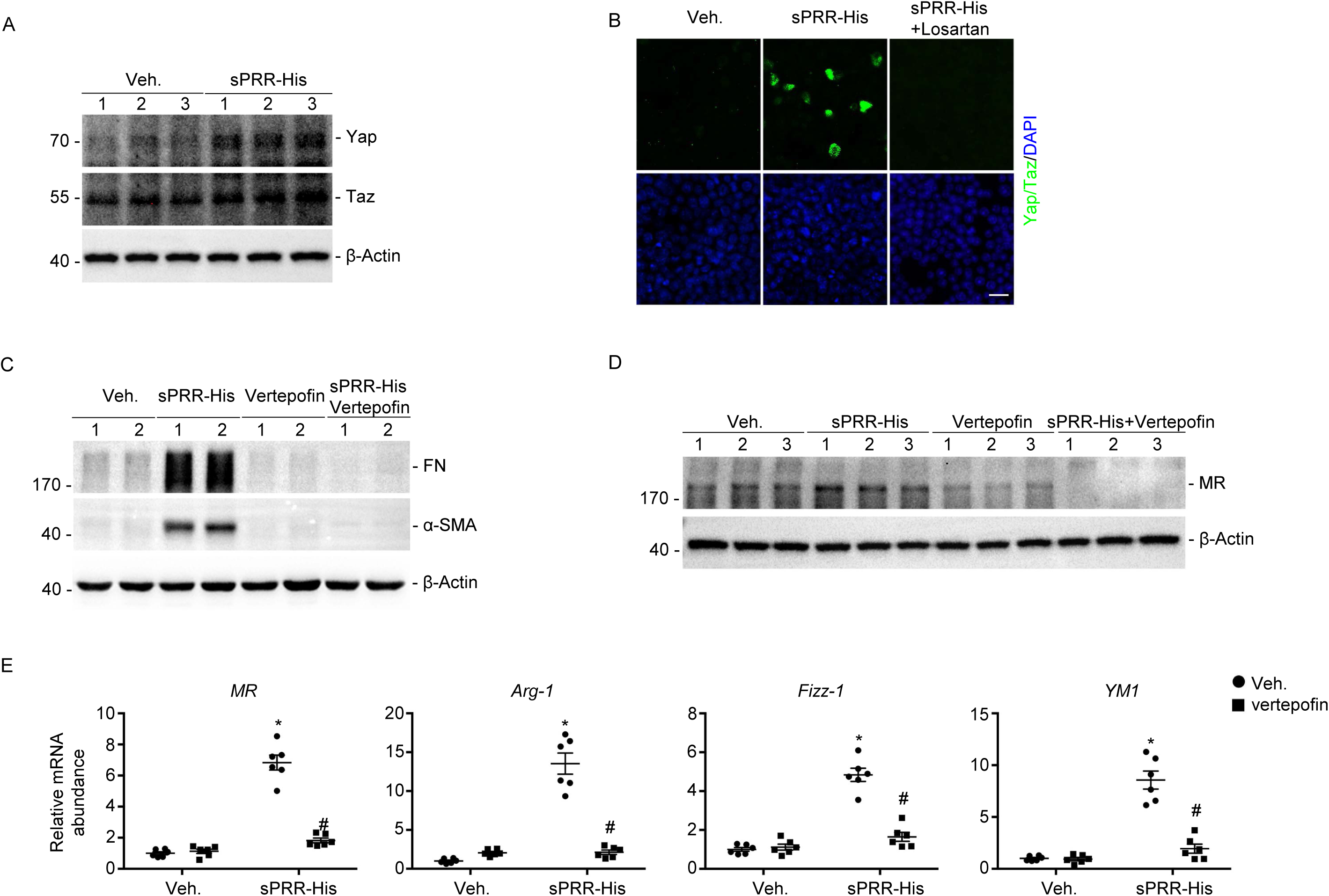
Yap/Taz mediates sPRR-His-induced macrophage alternative activation. (A) Western blot analyses showing abundance of Yap/Taz in BMDMs treated with 10 nM sPRR-His for 24 hours. Numbers (1-3) indicate individual sample within the same group. (B) Representative immunofluorescent staining images showing nuclear localization of Yap/Taz in BMDMs stimulated with sPRR-His or sPRR-His plus losartan. Scale bar = 20 μm. (C) Western blot analyses showing verteporfin inhibited sPRR-His-stimulated fibrosis in BMDMs. (D) Western blot analyses showing verteporfin inhibited sPRR-His-promoted macrophage alternative activation. (E) Real-time qRT-PCR analysis showing that verteporfin inhibited sPRR-His-stimulated macrophage alternative activation. BMDMs were pretreated with 10 nM verteporfin for 1 hour and then 10 nM sPRR-His for 24 hours. \**p <* 0.05 versus BMDMs treated with vehicle alone, n=6; ^#^*P*<0.05 versus BMDMs treated with sPRR-His, *n =* 6.

We studied the role of Yap/Taz induction in sPRR-His-induced macrophage M2 polarization and fibrosis, by treating BMDMs with the Yap inhibitor verteporfin,^30^ followed by sPRR-His treatment. The results showed that verteporfin notably inhibited the FN and α-SMA expression, cytosolic sPRR-His content, and MR expression stimulated by sPRR-His in BMDMs (Figures 7C and D). Additionally, verteporfin markedly decreased sPRR-His-induced mRNA expression of Arg-1, MR, Fizz-1, and YM1 in BMDMs (Figure 7E). These results indicated that sPRR-His may promote macrophage M2 polarization and fibrosis through activation of the Yap/Taz pathway.

## Discussion

We have recently reported that CD-specific deletion of PRR or suppression of sPRR production attenuates kidney fibrosis in a mouse model of UUO.^19^ As an extension of this observation, the present study addressed the underlying mechanisms of the profibrotic action of PRR/sPRR mediated by activation of macrophage Yap/Taz-dependent M2 polarization. We demonstrated that deletion of *PRR* in the CD or S1P inhibition attenuated macrophage M2 polarization accompanied by suppressed Yap/Taz expression in UUO mice. In cultured BMDMs, sPRR-His promoted macrophage M2 polarization associated with Yap/Taz in an AT1R-dependent manner.

Various types of renal disease undergo renal fibrosis, which leads to end-stage renal disease. Emerging evidence supports a profibrotic role of PRR. In cultured mesangial cells, PRR-mediated signaling leads to mesangial proliferation, type IV collagen accumulation, and increased expression of TGFβ1 mRNA through ERK1 and ERK2-dependent pathways.^31^ In cultured human proximal tubular cells, PRR mediates the indoxyl-sulfate-induced expression of TGFβ1 and α-SMA, which is expressed in myofibroblasts.^32^ Recently, we reported that sPRR promotes the fibrotic response in renal proximal tubule epithelial cells *in vitro* via the Akt/β-catenin/Snail signaling pathway.^33^ Additionally, PRR has been reported to regulate cell migration.^34^

The CD plays a key role in fine-tuning Na^+^ and water excretion and thus the homeostatic control of plasma volume and blood pressure. Surprisingly, increasing evidence supports a non-canonical role of the CD in the regulation of immunity and the pathogenesis of various types of renal disease. It has been reported that CD intercalated cells are involved in the recruitment of myeloid cells to the kidney, leading to AKI.^35^ We found that deletion of CD PRR attenuates macrophage accumulation in the obstructed kidneys. Within the injured kidneys, macrophages may be differentiated into different subtypes in response to various stimuli in the local microenvironment.^36^ M2 macrophages exhibit anti-inflammatory features and are involved in renal repair and fibrosis.^37^

We found that ablation of CD PRR inhibited macrophage M2 polarization in fibrotic kidneys. It has been shown that ablation of β-catenin in macrophages attenuates macrophage M2 polarization in fibrotic kidneys.^25^ PRR can promote kidney injury and fibrosis by amplifying Wnt/β-catenin signaling, which promotes macrophage alternative activation and kidney fibrosis.^38^ We found that ablation of CD PRR decreased macrophage M2 polarization and profibrotic cytokine expression. It seems likely that that CD PRR may regulate macrophage M2 polarization through β-catenin signaling.

The full-length PRR can be cleaved by furin or S1P and generate a 28-kDa sPRR.^26,39^ Plasma sPRR levels show a positive relationship with renal dysfunction.^40,41^ We found that ablation of CD PRR decreased the plasma sPRR level of mice subjected to 7-d UUO. We also confirmed that activation of CD PRR promoted macrophage accumulation, M2 polarization, and profibrotic cytokine expression via S1P-derived sPRR.

In cultured BMDMs, sPRR-His enhanced macrophage M2 polarization and expression of profibrotic cytokines. sPRR-His also partially inhibited the expression of macrophage M1 markers. Therefore, activation of CD PRR promotes kidney fibrosis by stimulating macrophage activation and M2 polarization through S1P-derived sPRR. M2 macrophage activation is driven by multiple cytokines and transcriptional factors.^22,42–44^ Activation of transcriptional regulators Yap/Taz mediates macrophage M2 polarization in UUO kidneys from mice,^23^ sPRR directly binds and activates AT1R in endothelial cells,^28^ and the YAP pathway is an important signaling branch downstream of AT1R.^24^

CD PRR/sPRR promotes macrophage M2 polarization by inducing the transcriptional regulators Yap/Taz based on the following reasons. Ablation of CD PRR resulted in less Yap/Taz expression and macrophage M2 polarization in UUO kidneys. Suppression of S1P-derived sPRR also markedly inhibited macrophage Yap/Taz induction and macrophage M2 polarization in UUO kidneys. sPRR-His up-regulates Yap/Taz expression in BMDMs. Down-regulating AT1R or Yap/Taz activation remarkably decreased sPRR-His-stimulated macrophage M2 polarization in BMDMs. TGFβ1, a well-known profibrotic cytokine, induces macrophage M2 polarization and Yap/Taz expression in cultured macrophages or fibroblasts.^23,45^

Furthermore, in BMDMs, inhibition of S1P markedly decreased TGFβ1-stimulated macrophage M2 polarization and Yap/Taz expression. Previous studies reported that Yap and Taz bind Smad 2/3 through the coiled-coil region, and this interaction may dictate the subcellular localization of Smad2/3.^46^ Further investigation is needed to explore the mechanisms for Yap/Taz-dependent regulation of macrophage M2 polarization.

## Conclusions

We have investigated the role and mechanisms of CD PRR in the pathogenesis of renal fibrosis in a UUO model. CD PRR promotes macrophage M2 polarization by regulating Yap/Taz activity and contributes to kidney fibrosis after UUO via S1P-derived sPRR as illustrated in the summary graph. Targeting PRR, sPRR, Yap, and Taz in the CD may be a new therapeutic strategy for protecting against kidney fibrosis in patients with CKD.

## NOVELTY AND RELEVANCE

### What Is New?

In our previous study, activation of the PRR/sPRR pathway in the distal nephron contributed to obstruction-induced renal fibrosis. However, the underlying mechanisms of this discovery have remained elusive. In the present study, we demonstrate that activation of the PRR/sPRR pathway promotes macrophage M2 polarization by regulating Yap/Taz activity, leading to kidney fibrosis in a unilateral ureteral obstruction model.

### What Is Relevant?

There is an unmet need to understand the molecular mechanisms and identify therapeutic targets to reverse renal fibrosis and halt the progression of chronic kidney disease to end-stage kidney disease. The present study contributes information about the novel role of the distal nephron in renal fibrosis.

### Clinical/Pathophysiological Implications?

Our findings support the concept that targeting the generation of sPRR in the distal nephron and its downstream signaling pathways may provide a potential therapy against renal fibrosis.

## Disclosure

None.

